# Long-read genome sequencing for the diagnosis of neurodevelopmental disorders

**DOI:** 10.1101/2020.07.02.185447

**Authors:** Susan M. Hiatt, James M.J. Lawlor, Lori H. Handley, Ryne C. Ramaker, Brianne B. Rogers, E. Christopher Partridge, Lori Beth Boston, Melissa Williams, Christopher B. Plott, Jerry Jenkins, David E. Gray, James M. Holt, Kevin M. Bowling, E. Martina Bebin, Jane Grimwood, Jeremy Schmutz, Gregory M. Cooper

## Abstract

**Purpose:** Exome and genome sequencing have proven to be effective tools for the diagnosis of neurodevelopmental disorders (NDDs), but large fractions of NDDs cannot be attributed to currently detectable genetic variation. This is likely, at least in part, a result of the fact that many genetic variants are difficult or impossible to detect through typical short-read sequencing approaches.

**Methods:** Here, we describe a genomic analysis using Pacific Biosciences circular consensus sequencing (CCS) reads, which are both long (>10 kb) and accurate (>99% bp accuracy). We used CCS on six proband-parent trios with NDDs that were unexplained despite extensive testing, including genome sequencing with short reads.

**Results:** We identified variants and created *de novo* assemblies in each trio, with global metrics indicating these data sets are more accurate and comprehensive than those provided by short-read data. In one proband, we identified a likely pathogenic (LP), *de novo* L1-mediated insertion in *CDKL5* that results in duplication of exon 3, leading to a frameshift. In a second proband, we identified multiple large *de novo* structural variants, including insertion-translocations affecting *DGKB* and *MLLT3*, which we show disrupt *MLLT3* transcript levels. We consider this extensive structural variation likely pathogenic.

**Conclusion:** The breadth and quality of variant detection, coupled to finding variants of clinical and research interest in two of six probands with unexplained NDDs strongly support the value of long-read genome sequencing for understanding rare disease.

## Introduction

Neurodevelopmental disorders (NDDS) are a heterogeneous group of conditions that lead to a range of physical and intellectual disabilities and collectively affect 1-3% of children^1^. Many NDDs result from large-effect genetic variation, which often occurs *de novo*^2^, with hundreds of genes known to associate with disease^3^. Owing to this combination of factors, exome and genome sequencing (ES/GS) have proven to be powerful tools for both clinical diagnostics and research on the genetic causes of NDDs. However, while discovery power and diagnostic yield of genomic testing have consistently improved over time^4^, most NDDs cannot be attributed to currently detectable genetic variation^5^.

There are a variety of hypotheses that might explain the fact that most NDDs cannot be traced to a causal genetic variant after ES/GS, including potential environmental causes and complex genetic effects driven by small-effect variants^6^. However, one likely possibility is that at least some NDDs result from highly penetrant variants that are missed by typical genomic testing. ES/GS are generally performed by generating millions of “short” sequencing reads, often paired-end 150 bp reads, followed by alignment of those reads to the human reference assembly and detection of variation from the reference. Various limitations of this process, such as confident alignment of variant reads to a unique genomic location, make it difficult to detect many variants, including some known to be highly penetrant contributors to disease. Examples of NDD-associated variation that might be missed include low-complexity repeat variants^7^, small to moderately-sized structural variants (SVs)^4,8^, and mobile element insertions (MEIs)^9,10^. Indeed, despite extensive effort from many groups, detection of such variation remains plagued by high error rates, both false positives and false negatives, and it is likely that many such variants are simply invisible to short read analysis^11^.

One potential approach to overcome variant detection limitations in ES/GS is to use sequencing platforms that provide longer reads, which allow for more comprehensive and accurate read alignment to the reference assembly, including within and near to repetitive regions, and *de novo* assembly^12^. However, to date, the utility of these long reads has been limited because of their high error rates. Recently, Pacific Biosciences released an approach, called Circular Consensus Sequencing (CCS), or “HiFi”, in which fragments of DNA are circularized and then sequenced repeatedly^13^. This leads to sequence reads that are both long (>10 kb) and accurate at the basepair level (>99%). In principle, such an approach holds great potential for more comprehensive and accurate detection of human genetic variation, especially in the context of rare genetic disease.

We have used CCS to analyze six proband-parent trios affected with NDDs that we previously sequenced using a typical Illumina genome sequencing (IGS) approach but in whom no causal or even potentially causal genetic variant, was found. The CCS data were used to detect variation within each trio and generate *de novo* genome assemblies, with a variety of metrics indicating that the results are more comprehensive and accurate, especially for complex variation, than those seen in short-read datasets. In one proband, we identified an L1-mediated *de novo* insertion within *CDKL5* that leads to a duplicated coding exon and is predicted to lead to a frameshift and loss-of-function. Transcript analyses confirm that the duplicated exon is spliced into mRNA in the proband. We have classified this variant as likely pathogenic using American College of Medical Genetics (ACMG) standards^14^. In a second proband, we found multiple large structural variants that together likely disrupt at least seven protein-coding genes. At a high level, these data strongly support the value of long-read genome analysis for the detection of NDD-associated variation, and more broadly for the analysis of human genetic disease.

## Materials and Methods

### Illumina sequencing, variant calling and analysis

Six probands and their unaffected parents were enrolled in a research study aimed at identifying genetic causes of NDDs^15^, which was monitored by Western IRB (20130675). All six of these families underwent trio Illumina genome sequencing (IGS) between four and five years ago, which was performed as described^15^. Briefly, whole blood genomic DNA was isolated using the QIAsymphony (Qiagen), and sequencing libraries were constructed by the HudsonAlpha Genomic Services Lab. Sequencing was performed on the Illumina HiSeqX using paired end reads with a read length of 150 base pairs. Each genome was sequenced at an approximate mean depth of 30X, with at least 80% of base positions reaching 20X coverage. While originally analyzed using hg37, for this study reads were aligned to hg38 using DRAGEN version 07.011.352.3.2.8b. Variants were discovered (in gvcf mode) with DRAGEN and joint genotyping was performed across six trios using GATK version 3.8-1-0-gf15c1c3ef. Structural variants (SVs) were called using a combination of Delly^16^, CNVnator^17^, ERDS^18^, and Manta^19^, followed by heuristic merging of SVs from the different callers based on breakpoint proximity and SV type. SVs are also annotated with gene features and allele frequencies from dbVar^20^, NDD publications^21,22^, and an internal SV database. Mobile element insertions (MEIs) were called using MELT^23^ run in MELT-SINGLE mode. Variant analysis and interpretation was performed using ACMG guidelines^14^, similar to that which we previously performed^4,15^. None of the probands had a Pathogenic (P), Likely Pathogenic (LP), or Variant of Uncertain Significance (VUS) identified by IGS, either at the time of original analysis or after a reanalysis performed at the time of generation of long-read data. In all trios, expected relatedness was confirmed^24^. IGS data for Probands 1-5 are available via dbGAP (https://www.ncbi.nlm.nih.gov/projects/gap/cgi-bin/study.cgi?study_id=phs001089.v3.p1). Project Accession Number: phs001089. Complete IGS data for proband 6 is not available due to consent restrictions.

### Long-Read sequencing, variant calling, analysis and de novo assemblies

Long-read sequencing was performed using Circular Consensus Sequence (CCS) mode on a PacBio Sequel II instrument (Pacific Biosciences of California, Inc.). Libraries were constructed using a SMRTbell Template Prep Kit 1.0 and tightly sized on a SageELF instrument (Sage Science, Beverly, MA, USA). Sequencing was performed using a 30 hour movie time with 2 hour pre-extension and the resulting raw data was processed using either the CCS3.4 or CCS4 algorithm, as the latter was released during the course of the study. Comparison of the number of high-quality indel events in a read versus the number of passes confirmed that these algorithms produced comparable results. Probands were sequenced to an average CCS depth of 30X (range 25 to 35), while parents were covered at an average depth of 16x (range 10 to 22, see Table 1, Supplemental Table 2A). CCS reads were aligned to the complete GRCh38.p13 human reference. For SNVs and indels, CCS reads were aligned using the Sentieon v.201808.07 implementation of the BWA-MEM aligner (https://www.biorxiv.org/content/10.1101/396325v1), and variants were called using DeepVariant v0.10^25^ and joint-genotyped using GLNexus v1.2.6 (https://www.biorxiv.org/content/10.1101/343970v1). For structural variants (SVs), reads were aligned using pbmm2 1.0.0 (https://github.com/PacificBiosciences/pbmm2) and SVs were called using pbsv v2.2.2 (https://github.com/PacificBiosciences/pbsv). Candidate *de novo* SVs required a proband genotype of 0/1 and parent genotypes of 0/0, with ≥ 6 alternate reads in the proband and 0 alternate reads, ≥5 reference reads in the parents. PacBio CCS data for Probands 1-5 will be submitted to dbGAP.

**Table 1.** Probands selected for PacBio sequencing.

| Family ID | Proband Gender | Race | Major Phenotypic Features | Previous Genetic Testing |  |  |  | PacBio CCS Coverage (P/D/M) | Average Insert Size(bp) (P/D/M) |
| --- | --- | --- | --- | --- | --- | --- | --- | --- | --- |
|  |  |  |  | Array | Single Gene Test(s) or Panel(s)<sup>a</sup> | ES/GS | Other Normal Test Results |  |  |
| 1 | F | C | Seizures, facial dysmorphism, hypotonia | Normal | Normal x2 | No Findings (both) | Karyotype | 25x/10x/11x | 12,655/12,238/12,884 |
| 2 | F | AA | ID, seizures, hypotonia | Normal | Normal x7 | No Findings (both) | Mito | 26x/16x/12x | 12,651/12,865/12,600 |
| 3 | M | C | ID, seizures | VUS dup | Normal x3 | No Findings (GS) | Fragile X | 35x/19x/22x | 14,393/16,604/16,344 |
| 4 | F | C/AA | ID, facial dysmorphism, hypotonia | Normal | Normal x1 | No Findings (GS) | Fragile X | 44x/14x/20x | 11,420/11,555/11,197 |
| 5 | M | C | ID, seizures, speech<br>delay, brain MRI<br>abnormalities | Normal | Normal x4 | No Findings<br>(GS) | Mito | 30x/16x/20x | 21,145/19,264/21,568 |
| 6 | F | C | ID, seizures, speech<br>delay | Normal | NP | No Findings<br>(GS) | NP | 33x/19x/14x | 12,452/12,183/13,641 |

ES/GS, exome sequencing/genome sequencing; P, proband; D, dad; M, mom; F, female; M, male; C, Caucasian; AA, African American; ID, intellectual disability; NP, not performed. <sup>a</sup> Some VUS SNVs have been reported in these probands.

For one proband (Proband 4), we used several strategies to create *de novo* assemblies using 44x CCS data. Assemblies were generated using canu (v1.8)^26^, Falcon unzip (falcon-kit 1.8.1-)^27^, HiCanu (hicanu_rc +325 changes (r9818 86bb2e221546c76437887d3a0ff5ab9546f85317))^28^, and hifiasm (v 0.5-dirty-r247)^29^. Hifiasm was used to create two assemblies. First, the default parameters were used, followed by two rounds of Racon(v1.4.10) polishing of contigs. Second, trio-binned assemblies were built using the same input CCS reads, in addition to kmers generated from a 36x paternal Illumina library and a 37x maternal Illumina library (singletons were excluded).The kmers were generated using yak(r55) using the suggested parameters for running a hifiasm trio assembly(kmer size=31 and Bloom filter size of 2**37). Maternal and paternal contigs went through two rounds of Racon(v1.4.10) polishing. Trio-binned assemblies were built for the remaining probands in the same way. Individual parent assemblies were also built with hifiasm (v0.5-dirty-r247) using default parameters. The resulting contigs went through two rounds of Racon(v1.4.10) polishing.

Coordinates of breakpoints were defined by a combination of assembly-assembly alignments using minimap2^30^ (followed by use of bedtools bamToBed), visual inspection of CCS read alignments, and BLAT. Rearranged segments in the chromosome 6 region were restricted to those >4 kb. Dot plots illustrating sequence differences were created using Gepard^31^.

### QC Statistics

SNV and indel concordance and *de novo* variant counts (Supplemental Tables 1A, 1C) were calculated using bcftools v1.9 and rtg-tools vcfeval v3.9.1. “High-quality *de novo”* variants were defined as PASS variants (IGS/GATK only) on autosomes (on primary contigs only) that were biallelic with DP ≥ 7 and genotype quality (GQ) ≥ 35. Additional requirements were a proband genotype of 0/1, with ≥ 2 alternate reads and an allele balance ≥ 0.3 and ≤ 0.7. Required parent genotypes were 0/0, with alternate allele depth of 0. Mendelian error rates were also calculated using bcftools (Supplemental Table 1D). “Rigorous” error rates were restricted to PASS variants (IGS/GATK only) on autosomes with GQ>20, and total allele depth (DP) >5. Total variant counts per trio (Supplemental Table 1B) were calculated using VEP (v98), counting multi-allelic sites as one variant. SV counts (Supplemental Table 1E) were calculated using bcftools and R. Counts were restricted to calls designated as “PASS”, with an alternate AD ≥2. Candidate SV *de novos* required proband genotype of 0/1 and parent genotypes of 0/0, with ≥ 6 alternate reads in the proband and 0 alternate reads, ≥5 reference reads in the parents. *De novo* MELT calls (Supplemental Table 5B) in IGS data were defined as isolated proband calls where the parent did not have the same type (ALU, L1, or SVA) of call within 1 kb as calculated by bedtools closest v.2.25.0. These calls were then filtered (using bcftools) for “PASS” calls and varying depths, defined as the number of read pairs supporting both sides of the breakpoint (LP, RP). To create a comparable set of *de novo* mobile element calls in CCS data (Supplemental Table 5D), individual calls were extracted from the pbsv joint-called VCF using bcftools and awk and isolated proband calls were defined as they were for the IGS data and filtered (using bcftools) for PASS calls and varying depths, defined as the proband alternate allele depth (AD[1]).

### Simple repeat and low mappability regions

We generated a bed file of disease-related low-complexity repeat regions in 35 genes from previous studies^7,32^. Most regions (25) include triplet nucleotide repeats, while the remainder include repeat units of 4-12 bp (Supplemental Table 4A). Reads aligning to these regions were extracted from bwa-mem-aligned bams and visualized using the Integrated Genomics Viewer (IGV^33^). Proband depths of MAPQ60 reads spanning each region (Supplemental Table 4A) were calculated using bedtools multicov v2.28.0. For the depth calculations, regions were expanded by 15 bp on either side (using bedtools slop) to count reads anchored into non-repeat sequence. The mean length of these regions was 83 bp, with a max of 133 bp.

Low mappability regions were defined as the regions of the genome that do not lie in Umap k100 mappable regions (https://bismap.hoffmanlab.org/)^34^. Regions ≥ 100,000 nt long and those on non-primary contigs were removed, leaving a total of 242,222 difficult-to-map regions with average length 411 bp. Proband depths of MAPQ60 reads spanning each region were calculated using bedtools multicov v2.28.0 (Supplemental Table 4B). High quality protein-altering variants (Supplemental Table 4C) in probands were defined using VEP annotations, and counted using bcftools v1.9. Requirements included a heterozygous or homozygous genotype in the proband, with ≥4 alternate reads, an allele balance ≥ 0.3 and ≤ 0.7, GQ>20, and DP>5. Reads supporting 57 loss-of-function variants (high-quality and low-quality) in Proband 5 were visualized with IGV and semi-quantitatively scored to assess call accuracy. Approximate counts of reads were recorded and grouped by mapping quality (MapQ=0 and MapQ≥1), along with subjective descriptions of the reads (Supplemental Table 4D). The total evidence across CCS and IGS reads was used to estimate truth and score each variant call as true positive (TP), false positive (FP), true negative (TN), false negative (FN), or undetermined (UN), see Supplemental Table 4D and 4E).

### CDKL5 cDNA Amplicon Sequencing

Total RNA was extracted from whole blood in PAXgene tubes using a PAXgene Blood RNA Kit version 2 (PreAnalytiX, #762164) according to the manufacturer’s protocol. cDNA was generated with a High Capacity Reverse Transcription Kit (Applied Biosystems, #4368814) using 500 ng of extracted RNA from each individual as input. Primers were designed to *CDKL5* exons 2, 5, and 6 to generate two amplicons spanning the potentially disrupted region of *CDKL5* mRNA. Select amplicons were purified and sent to MCLAB (Molecular Cloning Laboratories, South San Francisco, CA, USA) for Sanger sequencing. See Supplemental Methods for additional details, including primers.

### CDKL5 Genomic DNA PCR

We performed PCR to amplify products spanning both junctions of the insertion, in addition to the majority of the insertion using the genomic DNA (gDNA) of the proband and parents as template. Select amplicons were purified and sent to MCLAB (Molecular Cloning Laboratories, South San Francisco, CA, USA) for Sanger sequencing. See Supplemental Methods for additional details, including primers.

### DGKB/MLLT3 qPCR

Total RNA was extracted from whole blood using a PAXgene Blood RNA Kit version 2 (PreAnalytiX, #762164) and cDNA was generated with a High Capacity Reverse Transcription Kit (Applied Biosystems, #4368814) in an identical fashion as described for *CDKL5* cDNA amplicon sequencing. For qPCR, Two TaqMan probes targeting the MLLT3 exon 3-4 and exon 9-10 splice junctions (ThermoFisher, Hs00971092_m1 and Hs00971099_m1) were used with cDNA diluted 1:5 in dH2O to perform qPCR for six replicates per sample on an Applied Biosystems Quant Studio 6 Flex. Differences in CT values from the median CT values for either an unrelated family or the proband’s parents were used to compute relative expression levels. See Supplemental Methods for additional details, including primers.

## Results

Affected probands and their unaffected parents were enrolled in a research study aimed at identifying genetic causes of NDDs^15^. All trios were originally subject to standard Illumina genome sequencing (IGS) and analysis using ACMG standards^14^ to find pathogenic (P) or likely pathogenic (LP) variants, or variants of uncertain significance (VUS). Within the subset of probands for which no variants of interest (P, LP, VUS) were identified either originally or after subsequent reanalyses^4,15^, six trios were selected for sequencing using the PacBio Sequel II Circular Consensus Sequencing (CCS) approach (Table 1). These trios were selected for those with a strong suspicion of a genetic disorder, in addition to diversifying with respect to gender and ethnicity. Parents were sequenced, at a relatively reduced depth, to facilitate identification of *de novo* variation.

### QC of CCS data

Variant calls from CCS data and IGS data were largely concordant (Supplemental Table 1A). When comparing each individuals’ variant calls in the Genome in a Bottle (GIAB) high confidence regions^35^ between CCS and IGS, concordance was 94.63%, with higher concordance for SNVs (96.88%) than indels (75.96%). Concordance was slightly higher for probands only, likely due to the lower CCS read-depth coverage in parents. While CCS data showed a consistently lower number of SNV calls than IGS (mean = 7.0 M vs. 7.45 M, per trio), more *de novo* SNVs at high QC stringency were produced in CCS data than IGS (mean SNVs= 89 vs. 38, Supplemental Table 1B, 1C). CCS yielded far fewer *de novo* indels at these same thresholds (mean indels 11 vs. 148), with the IGS *de novo* indel count being much higher than biological expectation^36^ and likely mostly false positive calls (Supplemental Table 1C). In examining reads supporting variation that was uniquely called in each set, we found that CCS false positive *de novos* were usually false negative calls in the parent, due to lower genome-wide coverage in the parent and the effects of random sampling (i.e., sites at which there were 7 or more CCS reads in a parent that randomly happened to all derive from the same allele, Supplemental Table 1C). Mendelian error rates in autosomes were noticeably lower in CCS data relative to IGS (harmonic mean of high-quality calls 0.18% vs. 0.34%, Supplemental Table 1D), suggesting the CCS SNV calls are of higher accuracy, consistent with previously published data^13^.

Each trio had an average of ∼56,000 SVs among all three members, including an average of 59 candidate *de novo* SVs per proband (Supplemental Table 1E). Trio SVs mainly represent insertions (48%) and deletions (43%), followed by duplications (6%), single breakends (3%), and inversions (<1%).

Several assemblers were used to build *de novo* assemblies for one proband (Proband 4). Canu, Falcon, and HiCanu all produced high-quality assemblies, but hifiasm assemblies were of highest quality (Supplemental Table 2A). Use of trio-binned hifiasm allowed assembly of high quality maternal- and paternal-specific contigs with an average N50 of 45.65 Mb, approaching that of hg38. Trio-binned hifiasm *de novo* assemblies were also built for each proband. The average N50 for proband trio-based assemblies was 35.4 Mb (Supplemental Table 2B).

### Variation in Simple Repeat regions

Accurate genotyping of simple repeat regions like trinucleotide repeat expansions presents a challenge in short read data where the reads are often not long enough to span variant alleles. We assessed the ability of CCS to detect variation in these genomic regions, and compared that to IGS. We first examined variation in *FMR1* (MIM: 309550). Expansion of a trinucleotide repeat in the 5’ UTR of *FMR1* is associated with Fragile X syndrome (MIM: 300624), the second-most common genetic cause of intellectual disability^37^. Visualization of this region in all 18 individuals indicated insertions in all but two samples in the CGG repeat region of *FMR1* relative to hg38, with a range of insertion sizes from 6-105 bp (Supplemental Table 3, Supplemental Figure 1). When manually inspecting these regions, while one or two major alternative alleles are clearly visible, there are often minor discrepancies in insertion lengths, often by multiples of 3. It is unclear if this represents true somatic variation, or if this represents inaccuracy of consensus generation in CCS processing.

Like that for *FMR1*, manual curation of 34 other disease-causal repeat regions in each proband indicated that alignment of CCS reads provides a more accurate assessment of variation in these regions compared to IGS. When looking at region-spanning reads with high quality alignment (mapQ=60), 97% (34 of 35) of the regions were covered by at least 10 CCS reads in all six probands, as compared to 11% (4 of 35) of regions with high-quality IGS reads (Supplemental Table 4A). While all query regions measured ≤ 144 bp (which includes an extension of 15 bp on either end of the repeat region), seven query regions were ≥100 bp. When considering only regions of interest <100 bp, 14% (4 of 28 regions) are covered by at least 10 high-quality IGS reads in each proband. Mean coverage of high-quality, region-spanning reads across probands was higher in CCS data than in IGS (29 vs. 11, Supplemental Table 4A). Of all repeat regions studied, none harbored variation classified as P/LP/VUS.

We also compared coverage of high-quality CCS and IGS reads in low mappability regions of the genome, specifically those that cannot be uniquely mapped by 100 bp kmers^34^. While over half of these regions (62.5%) were fully covered by at least 10 high quality CCS reads (mapQ=60) in all six probands, only 19.3% of the regions met the same coverage metrics in the IGS data (Supplemental Table 4B). The average CCS read depth in these regions was 26 reads, vs. 8 reads in IGS. Within these regions, CCS yielded twice as many high quality, protein-altering variants in each proband when compared to IGS (182 in CCS vs. 85 in IGS) (Supplemental Table 4C). Outside of the low mappability regions, counts of protein-altering variants were similar (6,627 in CCS vs. 6,759 in IGS).

To assess the accuracy of the protein-altering variant calls in low-mappability regions, we visualized reads for 57 loss-of-function variants detected by CCS, IGS, or both in Proband 5 and used the totality of read evidence to score each variant as TP, FP, TN, FN, or undetermined. Six of these were “high-quality” calls (see Methods), and all of these were correctly called in CCS (TPs, 100%); in IGS, two were correctly called (TPs, 33%) and four were undetected (FNs, 67%) (Supplemental Table 4D). Among all 57 unfiltered variant calls, most CCS calls were correct (29 TP, 15 TN, total 77%) while most IGS calls were incorrect (16 FP, 22 FN, total 67%) (Supplemental Table 4E).

### Mobile Element Insertions

We searched for mobile element insertions (MEIs) in these six probands within the IGS data using MELT (Supplemental Table 5A, 5B)^23^ and within CCS data using pbsv (see Methods, Supplemental Table 1E, 5C, 5D). Our results suggest that CCS detection of MEIs is far more accurate. For example, it has been estimated that there exists a *de novo* Alu insertion in ∼1 in every 20 live births (mean of 0.05 per individual)^38,39^. However, at stringent QC filters (i.e., ≥5 read-pairs at both breakpoints, PASS, and no parental calls of the same MEI type within 1kb), a total of 82 candidate *de novo* Alu insertions (average of 13.7) were called across the six probands using the IGS data (Supplemental Table 5B), a number far larger than that expected. Inspection of these calls indicated that most were *bona fide* heterozygous Alu insertions in the proband that were inherited but undetected in the parents. Filtering changes to improve sensitivity come at a cost of elevated false positive rates; for example, requiring only 2 supporting read pairs at each breakpoint leads to an average of ∼55 candidate *de novo* Alu insertions per proband (Supplemental Table 5B). In contrast, using the CCS data and stringent QC filters (≥5 alternate reads, PASS, and no parental calls within 1kb) we identified a total of only 6 candidate *de novo* Alu MEIs among the 6 probands (Supplemental Table 5D), an observation that is far closer to biological expectation. We retained 4 candidate *de novo* Alu MEIs after further inspection of genotype and parental reference read depth (Supplemental Table 1E). One of these 4 appears genuine, while the other three appear to be correctly called in the proband but missed in the parents owing to low read-depth such that the Alu insertion-bearing allele was not covered by any CCS reads (Supplemental Figure 2).

### A likely pathogenic de novo structural variant in CDKL5

Analysis of structural variant calls and visual inspection of CCS data in proband 6 indicated a *de novo* structural variant within the *CDKL5* gene (MIM: 300203, Figure 1A). Given the *de novo* status of this event, the association of *CDKL5* with early infantile epileptic encephalopathy 2 (EIEE2, MIM: 300672), and the overlap of disease with the proband’s phenotype, which includes intellectual disability, developmental delay, and seizures, we prioritized this event as the most compelling candidate variant in this proband.

**Figure 1.**
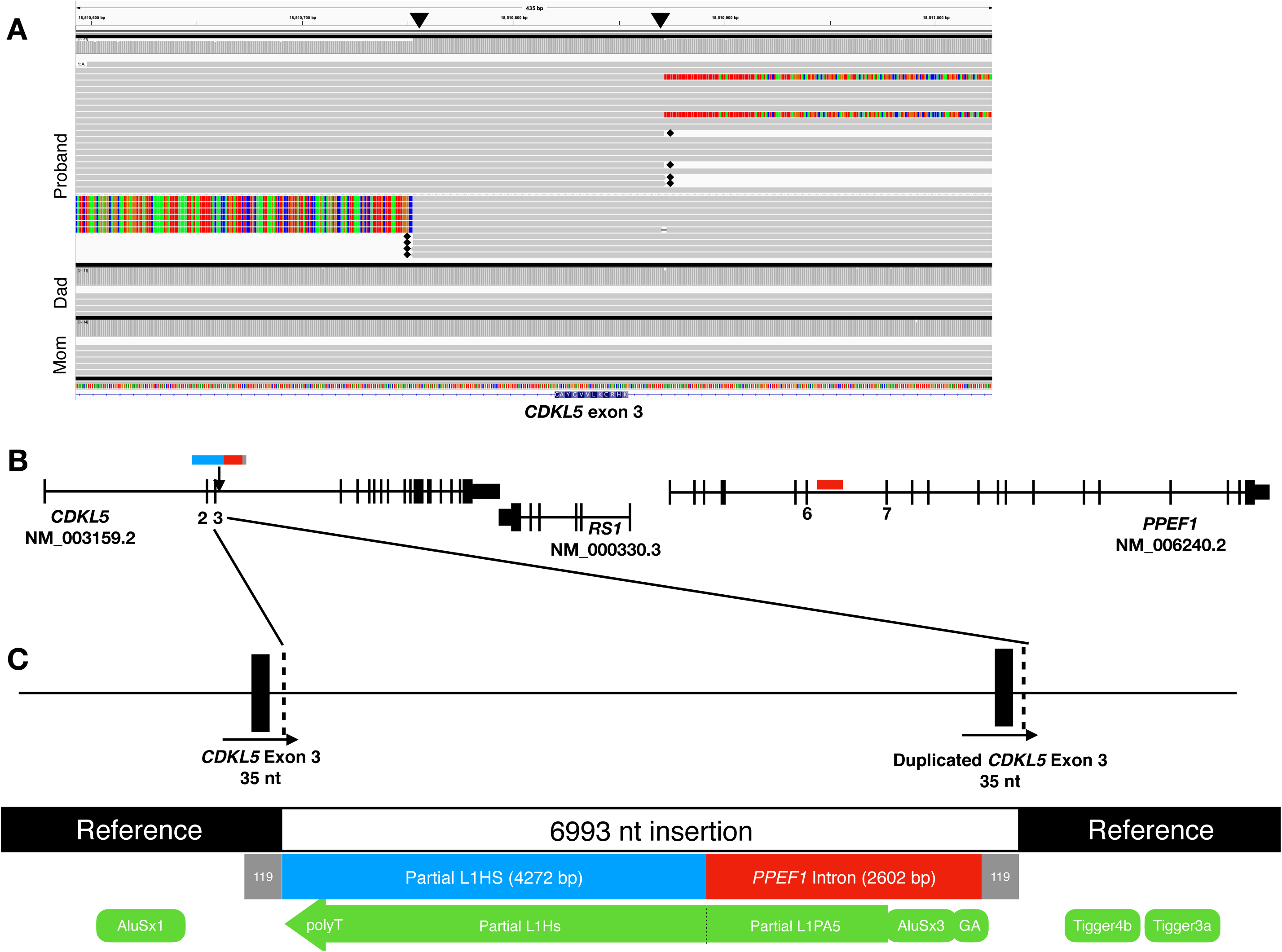
Proband 6 has a *de novo* insertion resulting in duplication of exon 3 of *CDKL5*. **A**. Alignment of CCS reads near exon 3 of *CDKL5* in IGV in Proband 6 and her parents. Unaligned portions of reads on either end of the 119 bp duplicated region are indicated with black triangles. The location of hard-clipped bases are designated with a black diamond. **B**. Gene structure of *CDKL5, RS1*, and *PPEF1*, indicating the location of the 6993 bp insertion in *CDKL5* and location of the duplicated *PPEF1* intronic sequence (red). **C**. Zoomed in view of the insertion. Black boxes indicate exons, gray boxes indicate the duplicated 119 bp segment, blue bar indicates a partial L1HS retrotransposon, and red indicates the duplicated *PPEF1* intronic sequence. Green boxes indicate RepeatMasker annotation of the proband’s insertion-bearing, contig sequence.

A trio-based *de novo* assembly in this proband identified a 45.3 Mb paternal contig and a 50.6 Mb maternal contig in the region surrounding *CDKL5*. While these contigs align linearly across the majority of the p arm of chromosome X (Supplemental Figure 3), alignment of the paternal contig to GRCh38 revealed a heterozygous 6993 bp insertion in an intron of *CDKL5* (GRCh38:chrX:18,510,871-18,510,872_ins6993, Figure 1, Supplemental Figure 4). Analysis of SNVs in the region surrounding the insertion confirm that it lies on the proband’s paternal allele. However, mosaicism is suspected, as there exist paternal haplotype reads within the proband that do not harbor the insertion (5 of 8 paternal reads without the insertion at the 5’ end of the event, and 7 of 16 paternal reads without insertion at the 3’ end of the event; Supplemental Figure 5).

Annotation of the insertion indicated that it contains three distinct segments: 4272 bp of a retrotransposed, 5’ truncated L1HS mobile element (including a polyA tail), 2602 bp of sequence identical to an intron of the nearby *PPEF1* gene (NC_000023.11:g.18738310_18740911; NM_006240.2:c.235+4502_235+7103), and a 119 bp region that includes a duplicated exon 3 of *CDKL5* (35 bp) and surrounding intronic sequence (GRCh38:chrX:18510753-18510871; NM_003159.2:c.65-67 to NM_003159.2:c.99+17; 119 bp total)(Figure 1B,C). The 2,602 bp copy of PPEF1 intronic sequence includes the 5’ end (1953 bp) of an L1PA5 element that is ∼6.5% divergent from its consensus L1, an AluSx element, and additional repetitive and non-repetitive intronic sequence. The size and identity of this insert in the proband, and absence in both parents, was confirmed by PCR amplification and Sanger sequencing (see Supplemental Methods).

Exon 3 of *CDKL5*, which lies within the target-site duplication of the L1-mediated insertion, is a coding exon that is 35 bp long; inclusion of a second copy of exon 3 into *CDKL5* mRNA is predicted to lead to a frameshift (Thr35ProfsTer52, Figure 2B). To determine the effect of this insertion on *CDKL5* transcripts, we performed RT-PCR from RNA isolated from each member of the trio. Using primers designed to span from exon 2 to exon 5, all three members of the trio had an expected amplicon of 240 bp. However, the proband had an additional amplicon of 275 bp (Figure 2A). Sanger sequencing of this amplicon indicated that a duplicate exon 3 was spliced into this transcript (Figure 2B). The presence of transcripts with a second copy of exon 3 strongly supports the hypothesis that the variant leads to a *CDKL5* loss-of-function effect in the proband.

**Figure 2.**
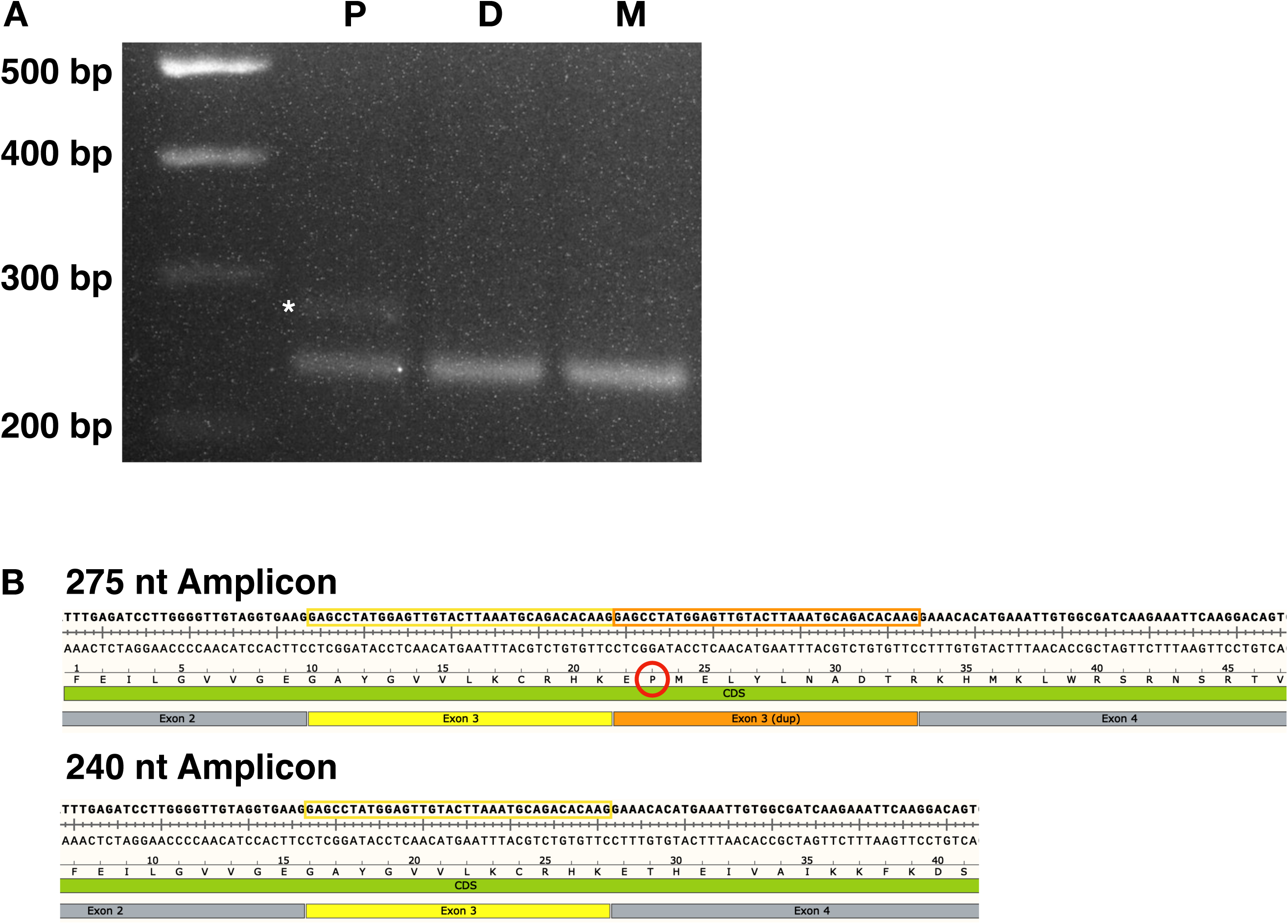
The duplicated *CDKL5* exon 3 is present in a subset of the proband’s *CDKL5* transcripts. **A**. RT-PCR using primers specific to exons 2-5 of *CDKL5* cDNA results in a 240 bp amplicon in proband (P), Dad (D), and Mom (M). An additional 275 bp amplicon is present only in the proband (asterisk). **B**. Sanger sequencing of both amplicons from the proband confirmed that the 240 bp amplicon includes the normal, expected sequencing and inclusion of a duplicated exon 3 in the upper, 275 bp band. This is predicted to lead to a frameshift (red circle) and downstream stop, p.(Thr35ProfsTer52). Yellow outlined box, exon 3 sequence; orange outlined box, duplicated exon 3 sequence.

### Multiple large de novo structural variants in Proband 4

Analysis of structural variant calls in proband 4 indicated several large, complex, *de novo* events affecting multiple chromosomes (6, 7, and 9). To elucidate the structure of the proband’s derived chromosomes, we inspected the trio-binned *de novo* assembly for this proband.

Four paternal contigs were assembled for chromosome 6, which showed many structural changes compared to reference chromosome 6 (Figure 3A). The proband harbors a pericentric inversion, with breakpoints at chr6:16,307,569 (6p22.3) and chr6:142,572,070 (6q24.2, Figure 3A, Supplemental Table 6A). In addition, a 9.3 Mb region near 6q22.31-6q23.3 contained at least eight additional breakpoints, with local rearrangement of eight segments, some of which are inverted (ABCDEFGH in reference vs. DCAGHFEB, Figure 3C, Table 6B). The median fragment size is just over 400 kb (range: 99 kb to 5.7 Mb, Supplemental Table 6B). While the ends of several fragments do overlap annotated repeats, many do not. We were not able to identify microhomology at the junctions of these eight segments. Together, the 10 breakpoints identified across chromosome 6 are predicted to disrupt at least six genes, five of which are annotated as protein-coding (Supplemental Table 6A). None of these have been associated with neurodevelopmental disease.

**Figure 3.**
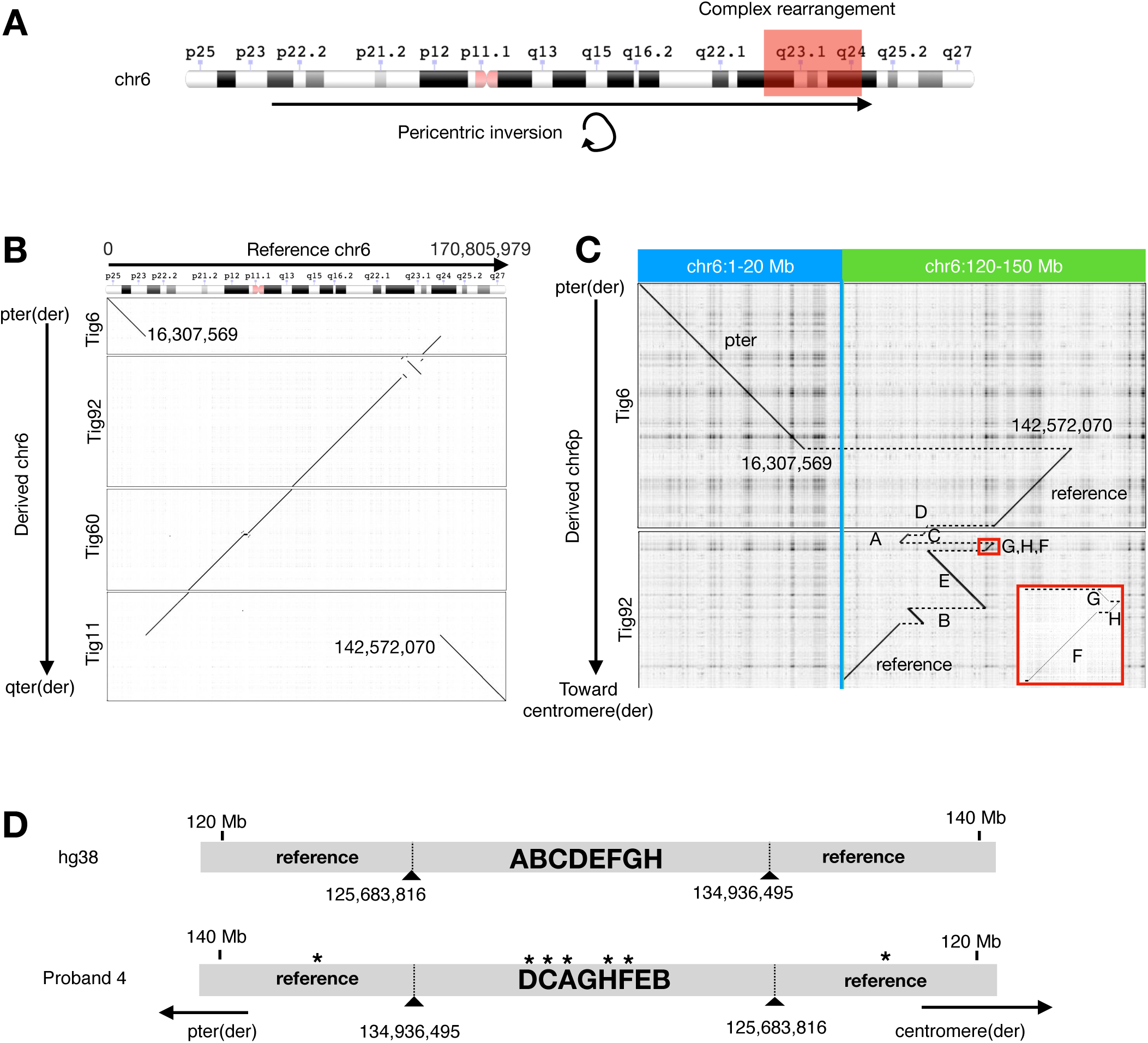
Proband 4 has several large structural changes on chromosome 6. **A**. Ideogram with annotation of structural variants on chromosome 6 identified in Proband 4. Ideogram is from the NCBI Genome Decoration Page (https://www.ncbi.nlm.nih.gov/genome/tools/gdp). **B**. Alignment of four sequential paternal contigs to reference chromosome 6 identified a pericentric inversion spanning 6p22.3 to 6q24.2 and a 9.3 Mb region near 6q22.31-6q23.3 with several additional breaks. **C**. Zoomed in view of 6q22.31-6q23.3 showing additional fragmentation. **D**. Schematic of complex rearrangement shown in C. Both reference hg38 structure and proband 4’s paternal allele structure are shown. Asterisks indicate inverted sequence as compared to hg38 reference.

CCS reads and contigs from the *de novo* paternal assembly of proband 4 also support structural variation involving chromosomes 7 and 9, with five breakpoints. The proband has two insertional translocations in addition to an inversion at the 5’ end of the chromosome 7 sequence within the derived 9p arm (Figure 4). Manual curation of SNVs surrounding all breakpoints confirmed that all variation lies on the paternal allele, and no mosaicism is suspected. Manual curation of the proband’s *de novo* assembly (specifically tig66) was required to resolve an assembly artifact (Supplemental Figure 6, Supplemental Methods).

**Figure 4.**
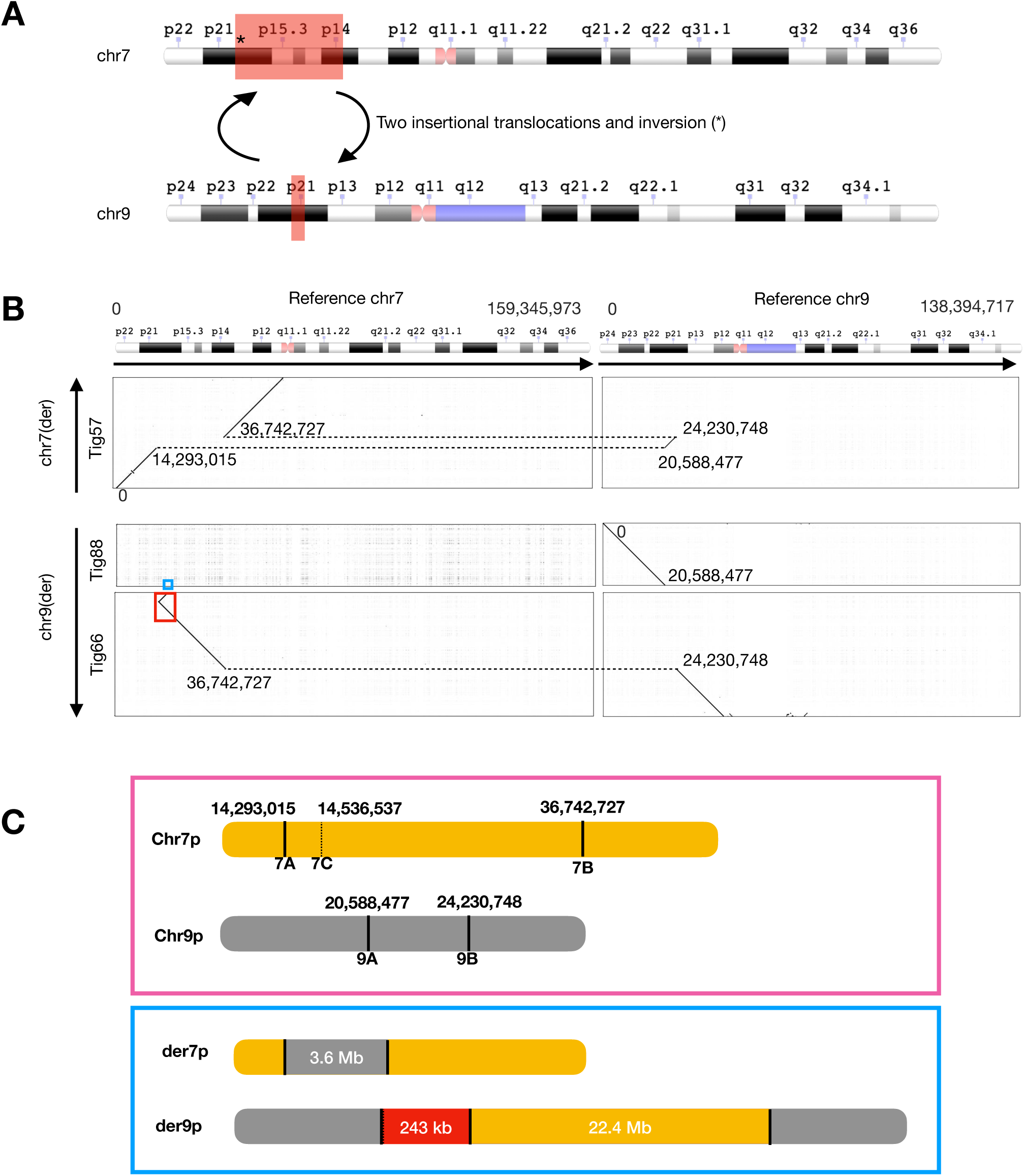
Proband 4 has two insertional translocations between chromosomes 7 and 9 and an inversion. **A**. Ideogram with annotation of rearrangements between chromosomes 7 and 9 identified in Proband 4. Ideograms are from the NCBI Genome Decoration Page (https://www.ncbi.nlm.nih.gov/genome/tools/gdp). **B**. Alignment of three paternal contigs to reference chromosomes 7 and 9 identified two insertional translocations. See Supplemental Figure 11 and Supplemental Methods regarding blue and red boxed areas. **C**. Schematic of the proband’s maternal (pink box) and paternal (blue box) p arms of chromosomes 7 and 9. The proband’s maternal alleles are annotated with corresponding hg38 coordinates. The paternal sequences represent the outcome of translocations (7A;9A and 7B;9B) and inversion (7A;7C), with fragment sizes shown. The red fragment in paternal der9p is inverted with respect to hg38 reference.

The net effect of the translocations and inversion is likely disruption of two protein-coding genes: *DGKB* (MIM: 604070) on chromosome 7 and *MLLT3* (MIM: 159558) on chromosome 9, neither of which have been associated with disease (Supplemental Table 6A). To determine if *MLLT3* transcripts are disrupted in this proband, we performed qPCR using RNA from each member of the trio, in addition to three unrelated individuals (Family 3). Using two validated TaqMan probes near the region of interest (exons 3-4 and exons 9-10), we found that proband 4 showed a ∼35-39% decrease in *MLLT3* compared to her parents and a 38-45% decrease relative to unrelated individuals (Supplemental Figure 7, Supplemental Table 7). Expression of *DGKB* was not examined, as the gene is not expressed at appreciable levels in blood^40^.

## Discussion

Here we describe CCS long-read sequencing of six probands with NDDs who had previously undergone extensive genetic testing with no variants found to be relevant to disease. Generally, the CCS genomes appeared to be highly comprehensive and accurate in terms of variant detection, facilitating detection of a diversity of variant types across many loci, including those that prove challenging to analysis with short reads. Detection of simple-repeat expansions and variants within low-mappability regions, for example, was far more accurate in CCS data than that seen in IGS, and many complex SVs were plainly visible in CCS data but missed by IGS.

Given the importance of *de novo* variation in rare disease diagnostics, especially for NDDs, it is also important to note the qualities of discrepant *de novo* calls between the two technologies. We found that most of the erroneously called *de novo* variants in the CCS data were correctly called as heterozygous in the proband but missed in the parents due to lower coverage and random sampling effects such that the variant allele was simply not covered by any reads in the transmitting parent. Such errors could be mitigated by sequencing parents more deeply. In contrast, *de novo* variants unique to IGS were enriched for systematic artifacts that cannot be corrected for with higher read-depth. Indels, for example, are a well-known source of error and heavily enriched among IGS de novo variant calls.

In one proband we identified a likely pathogenic, *de novo* L1-mediated insertion in *CDKL5. CDKL5* encodes cyclin-dependent kinase-like 5, a serine-threonine protein kinase that plays a role in neuronal morphology, possibly via regulation of microtubule dynamics^41^. Variation in *CDKL5* has been associated with EIEE2 (MIM: 300672), an X-linked dominant syndrome characterized by infantile spasms, early-onset intractable epilepsy, hypotonia, and variable additional Rett-like features^42,43^. *CDKL5* is one of the most commonly implicated genes identified by ES/GS in epilepsy cases^44^. Single nucleotide variants (SNVs), small insertions and deletions, copy-number variants (CNVs) and balanced translocations have all been identified in affected individuals, each supporting a haploinsufficiency model of disease^45^. We also note that *de novo* SVs, including deletions and at least one translocation, have been reported with a breakpoint in intron 3, near the breakpoint identified here^45–48^ (Supplemental Table 8, Supplemental Figure 8).

The variant harbors two classic marks of an L1HS insertion, including the preferred L1 EN consensus cleavage site (5’-TTTT/G-3’), and a 119-bp target-site duplication (TSD) which, in this case, includes exon 3 of *CDKL5*. Although TSDs are often fewer than 50 bp long, TSDs up to 323 bp have been detected^49^. The variant appears to be a chimeric L1 insertion. The 3’ end of the insertion represents retrotransposition of an active L1HS mobile element, with a signature polyA tail. However, the 5’ portion of the L1 sequence has greater identity to an L1 sequence within an intron of *PPEF1*, which lies about 230 kb downstream of *CDKL5*. Additional non-L1 sequence at the 5’ end of the insertion is identical to an intronic segment of *PPEF1*. While transduction of sequences at the 3’ end of L1 sequence has been described^50^, the *PPEF1* intronic sequence here lies at the 5’ end of the L1. A chimeric insertion similar to that observed here has been described previously, and has been proposed to result from a combination of retrotransposition and a synthesis-dependent strand annealing (SDSA)-like mechanism^49^.

Using ACMG variant classification guidelines, we classified this variant as Likely Pathogenic. The variant was experimentally confirmed to result in frameshifted transcripts due to exon duplication, and was shown to be *de novo*, allowing for use of both the PVS1 (loss of function)^51^ and PM2 (*de novo*)^52^ evidence codes. Use of Likely Pathogenic, as opposed to Pathogenic, reflects the uncertainty resulting from the intrinsically unusual nature of the variant and its potential somatic mosaicism, in addition to the fact that its absence from population variant databases is not in principle a reliable indicator of true rarity. Identification of additional MEIs and other complex structural variants in other individuals will likely aid in disease interpretation by both facilitating more accurate allele frequency estimation and by improving interpretation guidelines.

More generally, MEIs have been previously described as a pathogenic mechanism of gene disruption, but the contribution to developmental disorders has been limited to a modest number of cases in a few studies, each of which report P/LP variation lying within coding exons^9,10^. However, the MEI observed here in *CDKL5* would likely be missed by exome sequencing as the breakpoints are intronic, and in fact was also missed in our previous short-read genome sequencing analysis^15^. Global analyses of MEIs, such as our assessment of *de novo* Alu insertion rates (Supplemental Table 5), also support the conclusion that MEI events are far more effectively detected within CCS data compared to that seen in short read genomes. We find it likely that long-read sequencing will uncover MEIs that disrupt gene function and lead to NDDs in many currently unexplained cases.

CCS data also led to the detection of multiple large, complex, *de novo* structural variants in proband 4, affecting at least three chromosomes. While balanced translocations and pericentric inversions have been reported in healthy individuals, it is notable that both are present in proband 4, and additional events on both chromosomes 6 and 7 were identified at or near one of the breakpoints of the large rearrangements. The local rearrangement of eight segments near 6q22.31-6q23.3 appears to represent chromothripsis, as the segments are localized, do not have microhomology at their breaks, and show no significant copy gain or loss in the region (Supplemental Figure 9)^53^. The location of this cluster near one of the breakpoints of the pericentric inversion is consistent with observations that missegregated chromosomes can undergo micronucleus formation and shattering^54^. However, we cannot rule out other related mechanisms under the umbrella term of chromoanagenesis^55^.

Complex chromosomal rearrangements can lead to gene disruption and have been reported in individuals with NDDs or other congenital anomalies^56–58^. One of the most compelling disease causal candidate genes affected in proband 4 is *MLLT3*, which is predicted to be moderately intolerant to loss-of-function variation (pLI = 1, o/e = 0 (0 - 0.13)^59^; RVIS = 21.1%^60^). *MLLT3*, also known as *AF9*, undergoes somatic translocation with the *MLL* gene, also known as *KMT2A* (MIM: 159555), in patients with acute leukemia; pathogenicity in these cases results from expression of an in-frame KMT2A-MLLT3 fusion protein and subsequent deregulation of target HOX genes^61^. Balanced translocations between chromosome 4 and chromosome 9, resulting in disruption of *MLLT3*, have been previously reported in two individuals, each with NDDs including intractable seizures^62,63^. Although proband 4 does not exhibit seizures, she does have features that overlap the described probands, including speech delay, hypotonia, and fifth-finger clinodactyly.

While we cannot be certain of the pathogenic contribution of any one SV in proband 4, we consider the number, size, and extent of *de novo* structural variation to be likely pathogenic. ACMG recommendations on the interpretation of copy number variation were recently published, and although the events in proband 4 appear to be copy neutral, we attempted to apply modifications of these guidelines to these events^64^. The most compelling evidence for pathogenicity of these events is their *de novo* status (evidence code 5A), disruption of at least six protein-coding genes at the breakpoints (3A), at least one of which is predicted to be haploinsufficient (2H), and the total number of large SVs. While several of these can be captured by current evidence codes, they are weakened by the lack of affected disease-associated genes and the lack of a highly-specific phenotype in the proband. Further, although the SVs are large events, including a shattering of a >9 Mb region of the genome, we do not know the molecular effect on genes that are nearby but not spanning the breakpoints. Identification of additional complex structural variation like that in this proband will aid in development of additional guidelines for classification of these events.

Here we describe an analysis of six NDD-affected probands using PacBio CCS. These data facilitate more comprehensive and accurate detection of variation across a spectrum of categories, including low-complexity repeats, mobile element insertions, and complex structural variation. Among these newly detected variants, we identified likely pathogenic variation in two probands. While the sample size is too small to facilitate precise estimates of future yields in NDDs, two compelling positives from only six probands is consistent with the hypothesis that the ultimate yield among previously tested, but unsolved NDDs is substantial. This is likely also true for individuals suspected to have rare congenital disease more generally. Further, as CCS can capture complex variation in addition to essentially all variation detectable by short-read sequencing, it is likely that it will become a powerful front-line tool for research and clinical testing within rare disease genetics.

## Acknowledgements

This work was supported by a grant from The National Human Genome Research Institute, UM1HG007301. Some reagents were provided by PacBio as part of an early-access testing program. We thank our colleagues at HudsonAlpha who provided advice and general support, including Amy Nesmith Cox, Greg Barsh, Kelly East, Whitley Kelley, David Bick, and Elaine Lyon, in addition to the HudsonAlpha Genomic Services Laboratory and Clinical Services Laboratory. We also thank the clinical team at North Alabama Children’s Specialists. Finally, we are grateful to the families who participated in this study.

## Notes

**Conflicts of Interest** The authors all declare no conflicts of interest.

### Competing Interest Statement

The authors have declared no competing interest.

### Summary of Updates

Updated HTML abstract, text, Supplemental data, and Figures 3 and 4 to further describe variation in Proband 4.

