## Supplemental_Material for "Long-read genome sequencing for the diagnosis of neurodevelopmental disorders"

**Supplemental Methods**

*CDKL5 cDNA Amplicon Sequencing*

Amplicons were generated with CDKL5_Exon_2_Forward and either CDKL5_Exon_5_Reverse or CDKL5_Exon_6_Reverse. PCRs were performed with OneTaq 2x MM (NEB, M0482) with 10uM primers and 1uL cDNA input. Thermocycler conditions were as follows: 94$^{o}$C for 30 s, 40 cycles of 98$^{o}$C for 30 s, 56.4$^{o}$C for 30 s, 68$^{o}$C for 30 s, and a final extension of 68$^{o}$C for 5 min. To achieve full resolution of each amplicon, each PCR reaction was loaded onto a 2% agarose gel and ran at 120mV/hr for 2 hours prior to imaging. The CDKL5_Exon_5_Reverse primer containing reactions resulted in an expected ~240bp amplicon was observed for each sample in addition to a ~275bp amplicon uniquely observed in the proband. Similarly, the CDKL5_Exon_6_Reverse primer containing reactions resulted in an expected ~360bp amplicon for each sample in addition to a ~390bp amplicon uniquely observed in the proband. Each amplicon was gel extracted with a DNA gel recovery kit (Zymogen, #D4007) according to manufacturer's protocol and eluted in 20uL of pre-warmed elution buffer. To ensure sufficient input for Sanger sequencing, a second round of PCR that was identical the first round was performed with 1uL of gel extracted DNA as input from each amplicon. The products of this second round of PCR were again gel extracted and eluted in 20uL of pre-warmed elution buffer. Each elution was submitted to MCLAB (www.mclab.com) for Sanger sequencing.

The forward and reverse primer sequences were as follows:

CDKL5_Exon_2_Forward (NM_003159 cDNA location 222-245):

TGTGGCTTGCATCAAAAGAGGAGT

CDKL5_Exon_5_Reverse (NM_003159 cDNA location 441-462): TCCTGCTTGAGAGTCCGAAGCA

CDKL5_Exon_6_Reverse (NM_003159 cDNA location 561-582):

TCAGGTGGAACTCCATTTGGCA

*CDKL5 Genomic DNA PCR*

We performed PCR to amplify a product spanning the downstream junction (at GRCh38/hg38 chrX:18510868) using the genomic DNA (gDNA) of the proband and parents as template, in two separate reactions. PCR was performed using Q5 polymerase (NEB #M0494S) with 100 ng gDNA template and 0.5 uM each primer, with 30 seconds initial denaturation at 98°C, 35 cycles of 10 seconds 98°C denaturation, 20 seconds 68°C annealing, 1 minute 72°C extension, and a final 2 minutes 72°C extension. One reaction utilized primer LONG1_FWD and primer JUNC1_REV, resulting in an amplicon of 729 bp, only with proband gDNA. The second reaction utilized primer LONG2_FWD and primer JUNC1_REV, resulting in an amplicon of 1,207 bp, only with proband gDNA. We performed agarose gel electrophoresis and observed no bands in these reactions with parents' gDNA as template, and the correct size bands with the proband gDNA as template. The amplicons were purified with QiaQuick columns (Qiagen #28104) and were Sanger sequenced (MCLab) using the same primers utilized in the PCR; the sequences were aligned to and matched the assembled proband sequence.

We performed a long PCR to amplify from the position of JUNC1_REV across the entirety of the insert sequence and into the upstream gDNA sequence of the proband. PCR was performed using Q5 polymerase (NEB #M0494S) with 100 ng gDNA template and 0.5 uM each primer, with 30 seconds initial denaturation at 98°C, 35 cycles of 10 seconds 98°C denaturation, 8 minutes

68°C annealing/extension, 8 minutes 72°C extension. We increased the time at 68°C primarily for polymerase extension across the A/T-rich sequence in the insert (Dhatterwal, et al., PMID: 29183338). The reaction utilized the primers LONG3_FWD; reverse complement of JUNC1_REV and LONG3_REV, resulting in an amplicon of 7,989 bp only with proband gDNA. We performed agarose gel electrophoresis and observed no bands in these reactions with parents' gDNA as template, and the correct size band with the proband gDNA as template. The amplicon was purified with a QiaQuick column (Qiagen #28104) and was Sanger sequenced (MCLab). For Sanger sequencing, we utilized seven primers: LONG3_FWD and LONG3_REV (the primers used to generate the amplicon), and LONGVAL1-LONGVAL5. This resulted in five non-contiguous regions of the amplicon, comprising in total approximately 57% of its length, to be sequence verified.

To verify PPEF1 sequence in the proband insert, we performed two paired PCR assays. PCR was performed using Q5 polymerase (NEB #M0494S) with 100 ng gDNA template and 0.5 uM each primer, with 30 seconds initial denaturation at 98°C, 35 cycles of 10 seconds 98°C denaturation, 20 seconds 68°C annealing, 1 minute 72°C extension, and a final 2 minutes 72°C extension. One assay utilized the primer JUNC3_REV (matches PPEF1 insert sequence) with the primer PPEF1_FWD2 (matches PPEF1 sequence not present in insert), resulting in an amplicon of 1,122 bp in proband and parental gDNA; this was paired with a PCR using the same JUNC3_REV and the primer JUNC3_FWD (matches CDKL5 sequence outside the insert), resulting in an amplicon of 1,074 bp in the proband gDNA but not in the parents' gDNA. The other assay utilized the primer JUNC4_REV (matches PPEF1 insert sequence) with the primer PPEF1_FWD3 (matches

PPEF1 sequence not present in insert), resulting in an amplicon of 1,004 bp in proband and parental gDNA; this was paired with a PCR using the same JUNC4_REV and the primer JUNC4_FWD (matches CDKL5 sequence outside the insert), resulting in an amplicon of 821 bp in the proband gDNA but not in the parents' gDNA. We performed agarose gel electrophoresis and observed the correctly sized bands (and appropriate absence of bands) in all these reactions.

Primers:

LONG1_FWD: 5'-AACCTGTACATGCCCACACG-3'

JUNC1_REV: 5'-GCCCCGTTGTGTCTGTTTTC-3'

LONG2_FWD: 5'-TATGAGGTGCACGGCATAGG-3'

JUNC1_REV: 5'-GCCCCGTTGTGTCTGTTTTC-3'

LONG3_FWD: 5'-GAAAACAGACACAACGGGGC-3'

LONG3_REV: 5'-acacacCCCTGTCAAGCAAA-3'

LONGVAL1: 5'-TCTCACGTGCAGAGACACAC-3'

LONGVAL2: 5'-GTGTGTCTCTGCACGTGAGA-3'

LONGVAL3: 5'-GAGGCCAGGAGTTTGAGACC-3'

LONGVAL4: 5'-CACCACCGATCCCACAGAAA-3'

LONGVAL5: 5'-TGAGGAATCGCCACACTGAC-3'

JUNC3_REV: 5'-ATCTGTCACGGCTTCCCTTG-3'

PPEF1_FWD2: CTTCCACCCACTCCCCATTC-3'

JUNC3_FWD: GTACATGCCCACACGCAAAG-3'

JUNC4_REV: GCAGGCAATGGAGGTGTAGT-3’

PPEF1_FWD3: 5’-agcagcagccagactcaaat-3’

JUNC4_FWD: 5’-TCATTATGAGGTGCACGGCA-3’

**Resolution of tig66 mis-assembly in Proband 4**

One of the paternal contigs in the proband’s *de novo* assembly (tig66) appeared to be misassembled, likely due to the chr7 sequence inversion. Manual inspection of the proband’s *de novo* assembly vs. hg38 assembly alignments (and vice versa) and SNVs in these regions showed the paternal assembly contained both paternal and maternal sequence. This is a documented artifact of the current version of hifiasm (https://github.com/chhylp123/hifiasm/issues/10; https://github.com/chhylp123/hifiasm/issues/21). Curation of SNVs in contigs and CCS reads allowed manual reassembly of this region. Visualization of the non-trio-binned hifiasm assembly also confirmed the proper alignments (Supplemental Figure 6).

**Supplemental Table 3.** Range of insertion sizes for *FMR1* 5' UTR variation compared to GRCh38, based on visualization of reads in IGV.

| Family ID | Individual | Gender | Allele 1, nt insertion | Allele 2, nt insertion |
|---|---|---|---|---|
| 1 | C | F | 75 | 63-66 |
| | D | M | NA | 63 |
| | M | F | 72 | 66 |
| 2 | C | F | 33 | 27 |
| | D | M | NA | 27 |
| | M | F | 33 | 6 |
| 3 | C | M | 30 | NA |
| | D | M | NA | 66 |
| | M | F | 27-30 | 27-30 |
| 4 | C | F | 39-42 | 33 |
| | D | M | NA | 33 |
| | M | F | 36-39 | 36-39 |
| 5 | C | M | 30 | NA |
| | D | M | 33 | NA |
| | M | F | 105 | 30 |
| 6 | C | F | 69 | 9 |

| | | | | |
|---|---|---|---|---|
| | D | M | NA | 9 |
| | M | F | 69 | 45 |

C, child; D, dad; M, mom; NA, not applicable for hemizygous males.

**Supplemental Table 8.** Pathogenic or likely pathogenic structural variants that have been reported near intron 3 of *CDKL5.*

| **Publication** | **Proband in Pub** | **detected by** | **GRCh38 Coordinates** | **Flanking sequence** | **Description** | **Inheritance** |
|---|---|---|---|---|---|---|
| This Study | 6 | CCS | chrX:18510871 | AluSx/Tigger3a/4b | LINE/*PPEF1* insertion and exon 3 dup | *de novo* |
| Erez, et al.[a] | 1 | array | chrX:18369553-18526954 | AluSq; AluSp | 157409 bp del, removes exon 1-3 | unknown |
| Erez, et al. [a] | 2 | array | chrX:18432728-18570428 | AluJb; unique | 137701 bp del; removed exons 1-4 | *de novo* |
| Erez, et al. [a] | 3 | array | chrX:18439733-18514587 | AluSx; AluSq | 74869 bp del; removes exons 1-3 | *de novo* |
| Bartnik et al.[b] | 1 | array | chrX:18406582-18521160 | L1MB5; unique | 114579 bp del; removes exons 1-3 | *de novo* |
| Bartnik et al. [b] | 2 | array | chrX:18564194-18564780 | unknown | exon 4 del (confirmed by RT-PCR) | *de novo* |

| | | | | | | |
|---|---|---|---|---|---|---|
| Cordova-Fletes, et al.[c] | 1 | karyotype; FISH; RT-PCR; array painting | chrX:18519860-18532094 | unknown | t(X;2)(p22.1;p25.3) | unknown |
| Sanchis-Juan, et al.[d] | 4 | WGS; ONT long reads | chrX:17774889-18055885 | Alu | 280 kb dup | *de novo* |
| Sanchis-Juan, et al.[d] | 4 | WGS; ONT long reads | chrX:17774889-18514192 | Alu | 458 kb inversion | *de novo* |
| Sanchis-Juan, et al.[d] | 4 | WGS; ONT long reads | chrX:18230835-18514192 | Alu | 283 kb dup | *de novo* |

[a] PMID:19471977, [b] PMID:21293276, [c] PMID:19807736, [d] PMID:30526634

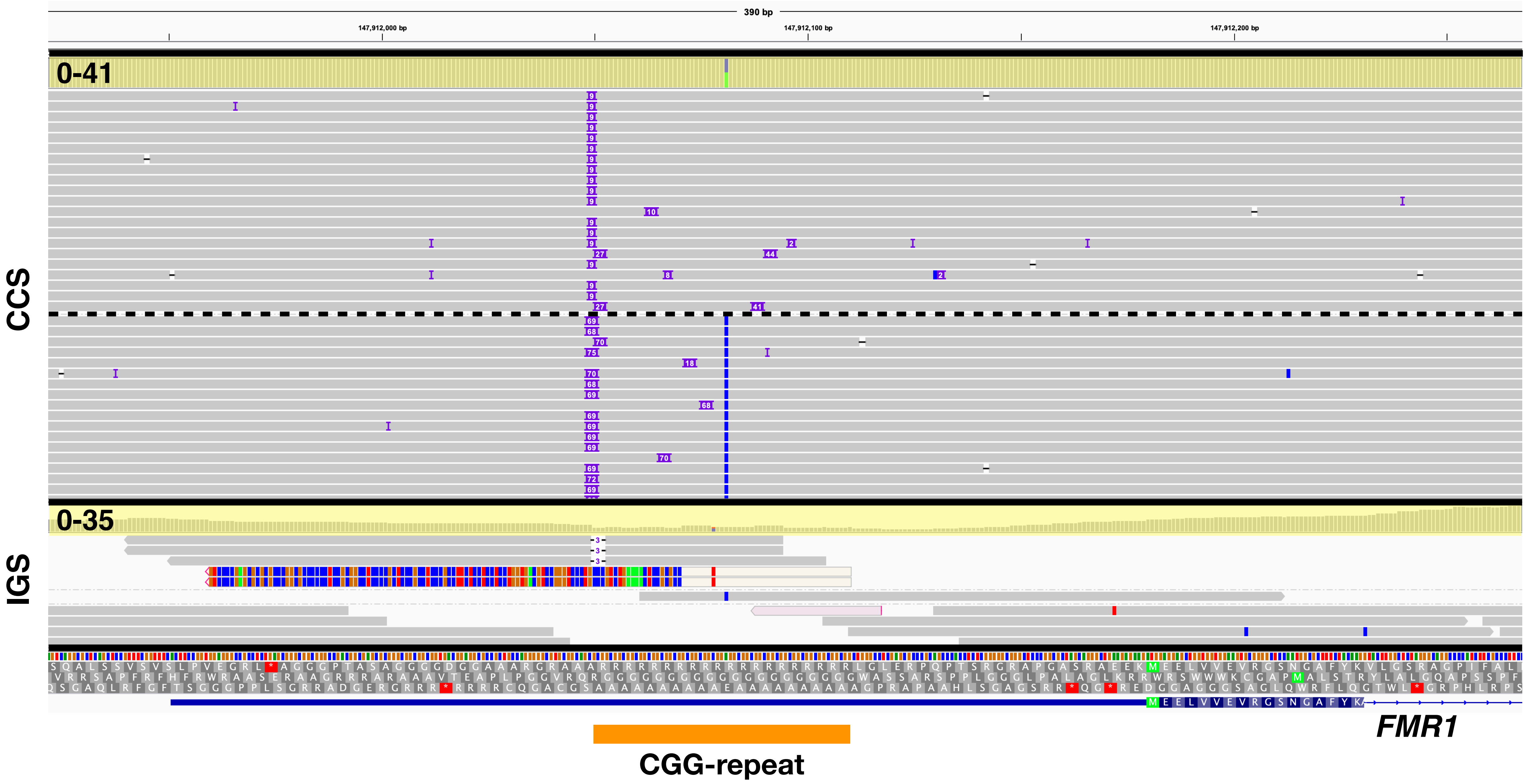


**Supplemental Figure 1.** Visualization of a subset of CCS and IGS reads in Proband 6 aligned to the 5’ end of FMR1 which contains a CGG-repeat region (orange bar). CCS reads group into two bins to represent two distinct repeat alleles (69 nt and 9 nt alleles), separated by a dashed line. Note read depth highlighted by the yellow box, with depth scale shown at the left end of the box.

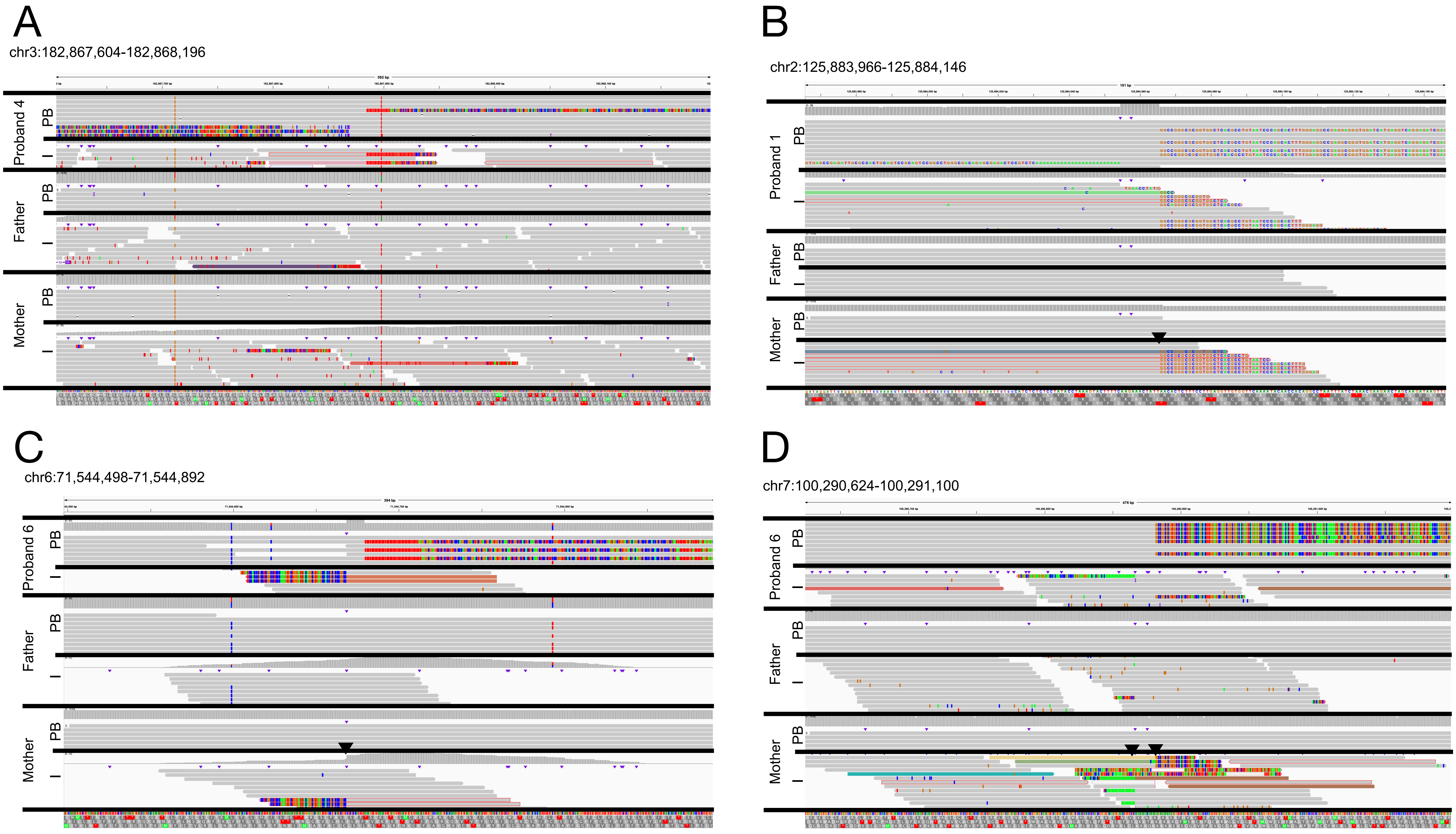


**Supplemental Figure 2.** Four *de novo* Alu insertions were called from CCS data in the six probands. In one case, CCS and IGS reads both appear to support *de novo* status (A), while in the three other cases (B, C, D), there are reads supporting the *Alu* insertion in the mom's Illumina reads (black triangle).

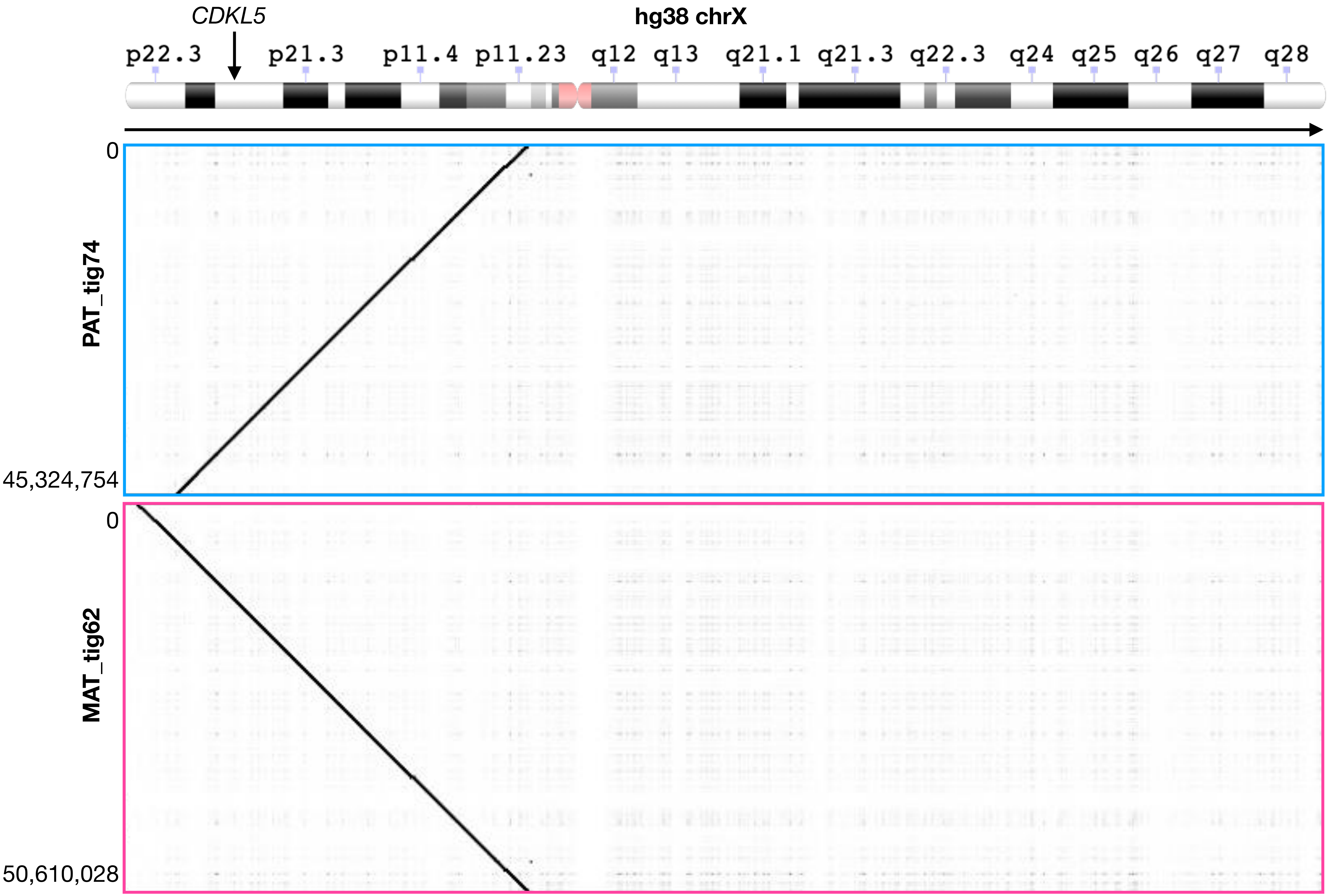


**Supplemental Figure 3.** *De novo* trio-based assembly for proband 6 resulted in two large paternal- and maternal-specific contigs (PAT_tig74 and MAT_tig62) covering the majority of the p arm of chromosome X, including *CDKL5*. Ideogram is from the NCBI Genome Decoration Page (https://www.ncbi.nlm.nih.gov/genome/tools/gdp).

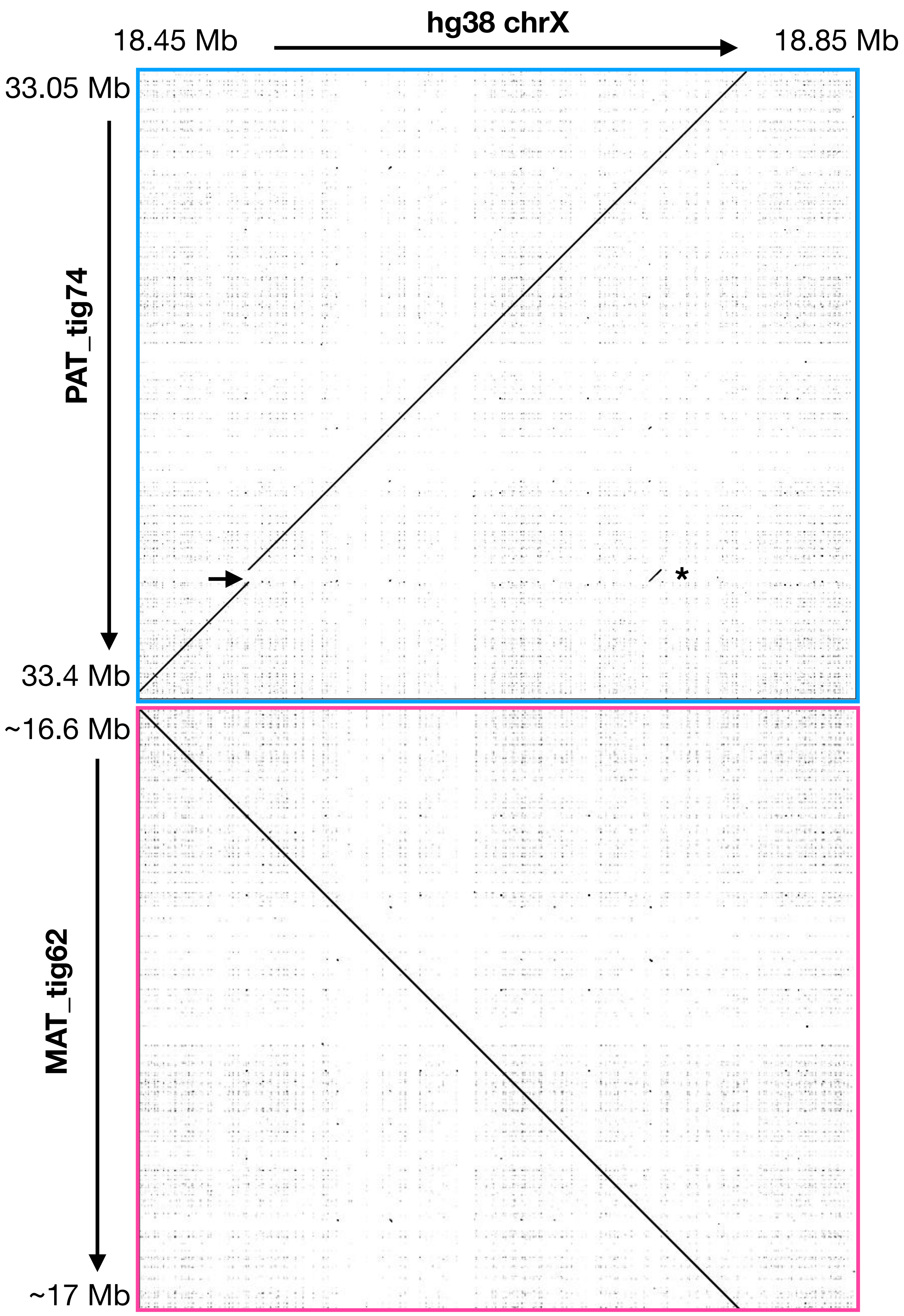


**Supplemental Figure 4.** Alignment of Proband 6's paternal and maternal contigs surrounding *CDKL5* to reference chromosome X. The 6993 bp insertion in the proband's primary contig is noted with an arrow, and alignment to a downstream LINE in an intron of *PPEF1* is shown with an asterisk.

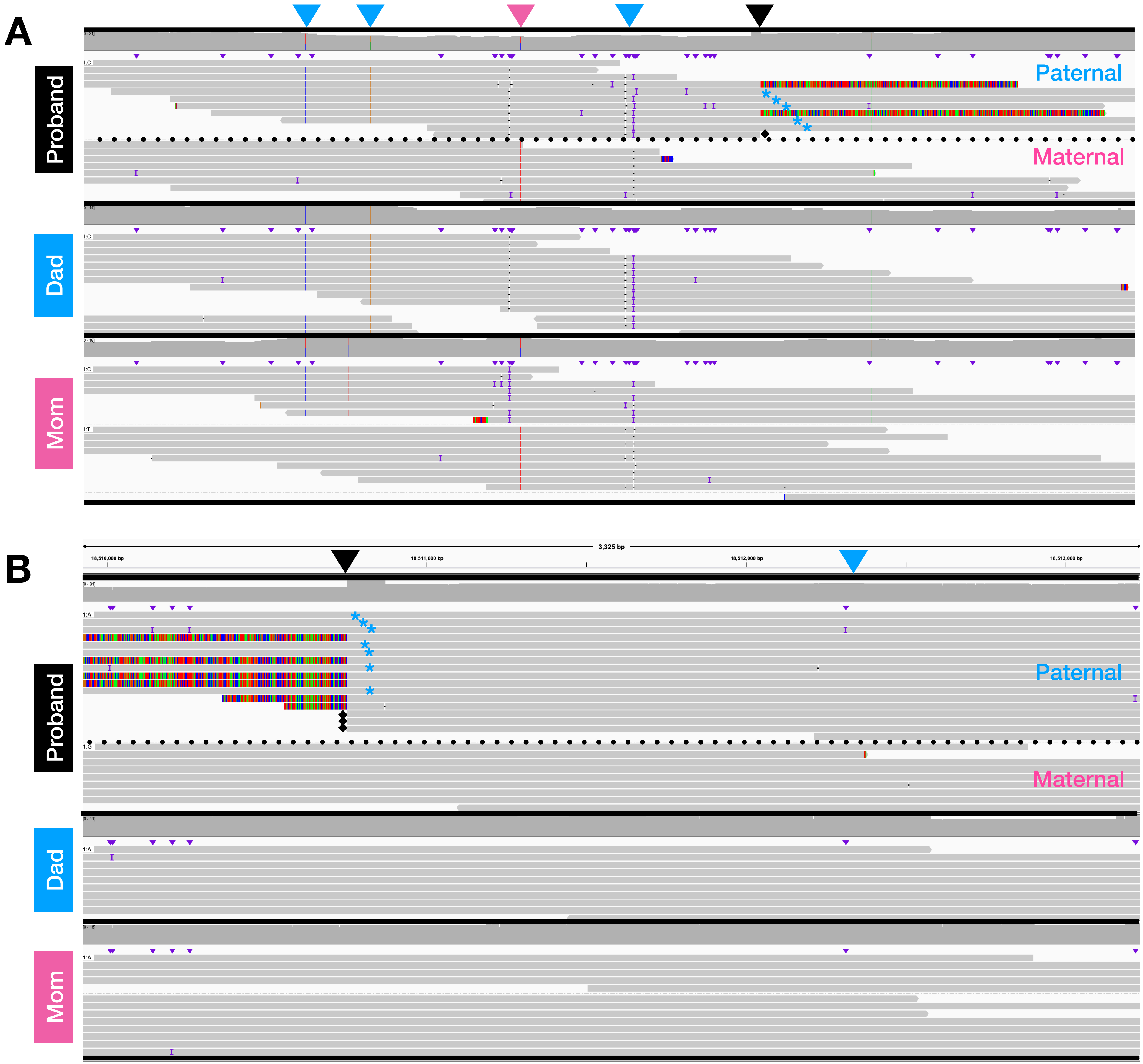


**Supplemental Figure 5.** Phasing of single nucleotide variants both upstream (A) and downstream (B) of the insertion (black triangle) in *CDKL5* in proband 6 indicate that the insertion is on the paternal allele. Inherited variants on the paternal allele (blue triangles) and maternal allele (pink triangle) are labeled. A dotted line separates the proband's haplotypes, which are labeled as paternal or maternal. Note the existence of paternal alleles lacking the insertion, suggesting mosaicism (blue asterisks). Black diamonds indicate reads that are hard-clipped and support the insertion. All reads supporting the insertion are shown, but only a subset of other reads are shown here.

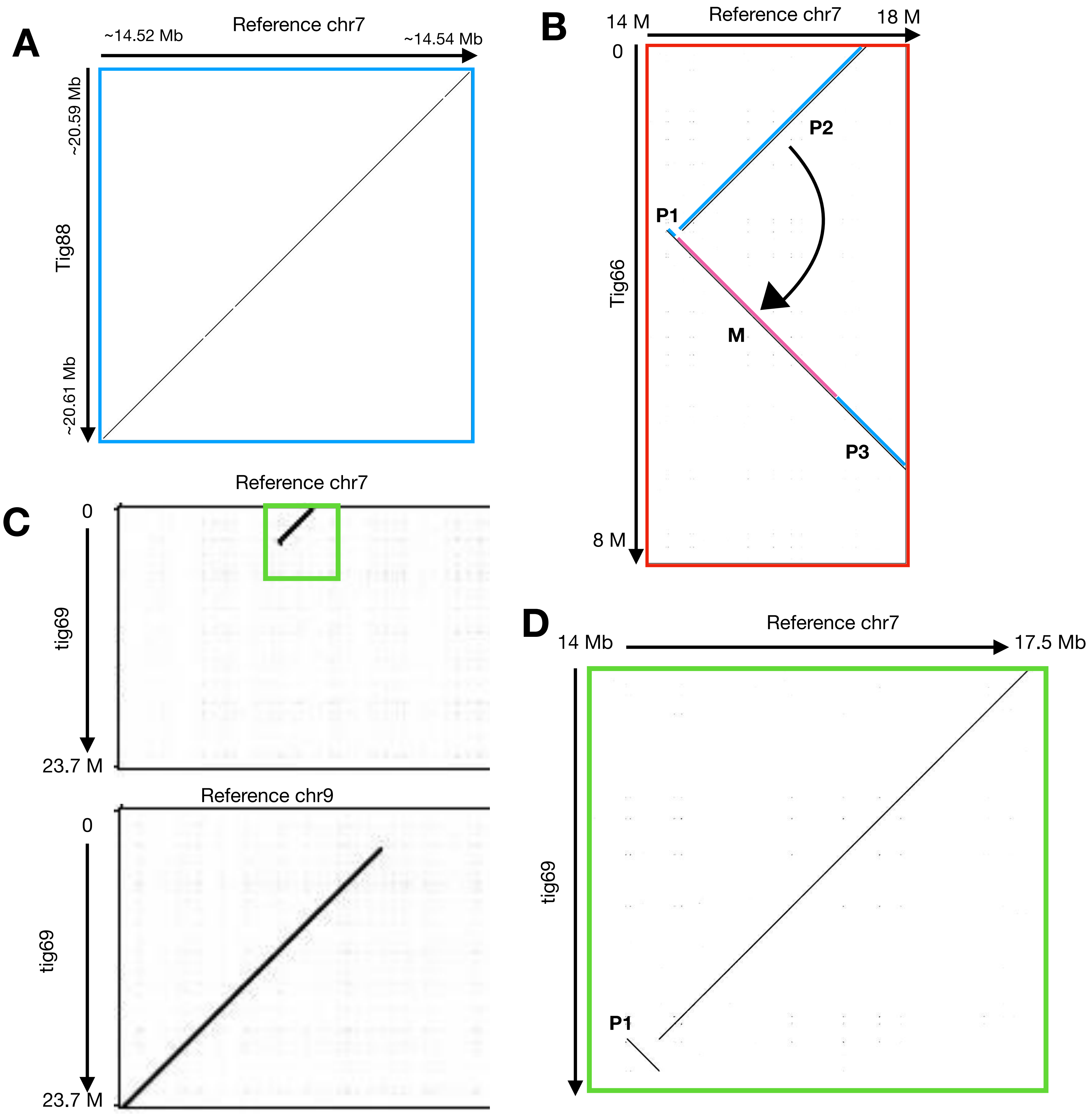


**Supplemental Figure 6.** Manual curation of an assembly artifact was required for PAT_tig66. **A.** Zoomed in view of blue box from Main Figure 4B. This region shows the alignment at the 3' end of tig88 to chromosome 7, and the sequence is inverted compared to the neighboring, upstream chr9 sequence. **B.** Zoomed in view of red box from Figure 4B. Manual inspection of this paternally-binned contig revealed both maternal and paternal sequence. P1, P2 and P3 represent paternal sequences, M represents maternal sequence. If the maternal sequence is removed, and P2 is aligned contiguously with P3, P1 represents an inversion with respect to these sequences. **C.** A contig (tig69) from same region from the non-trio-binned hifiasm assembly does not contain this error. **D** Zoomed in region from C. Tig69 aligned to chromosome 7 sequence shows the inversion (P1).

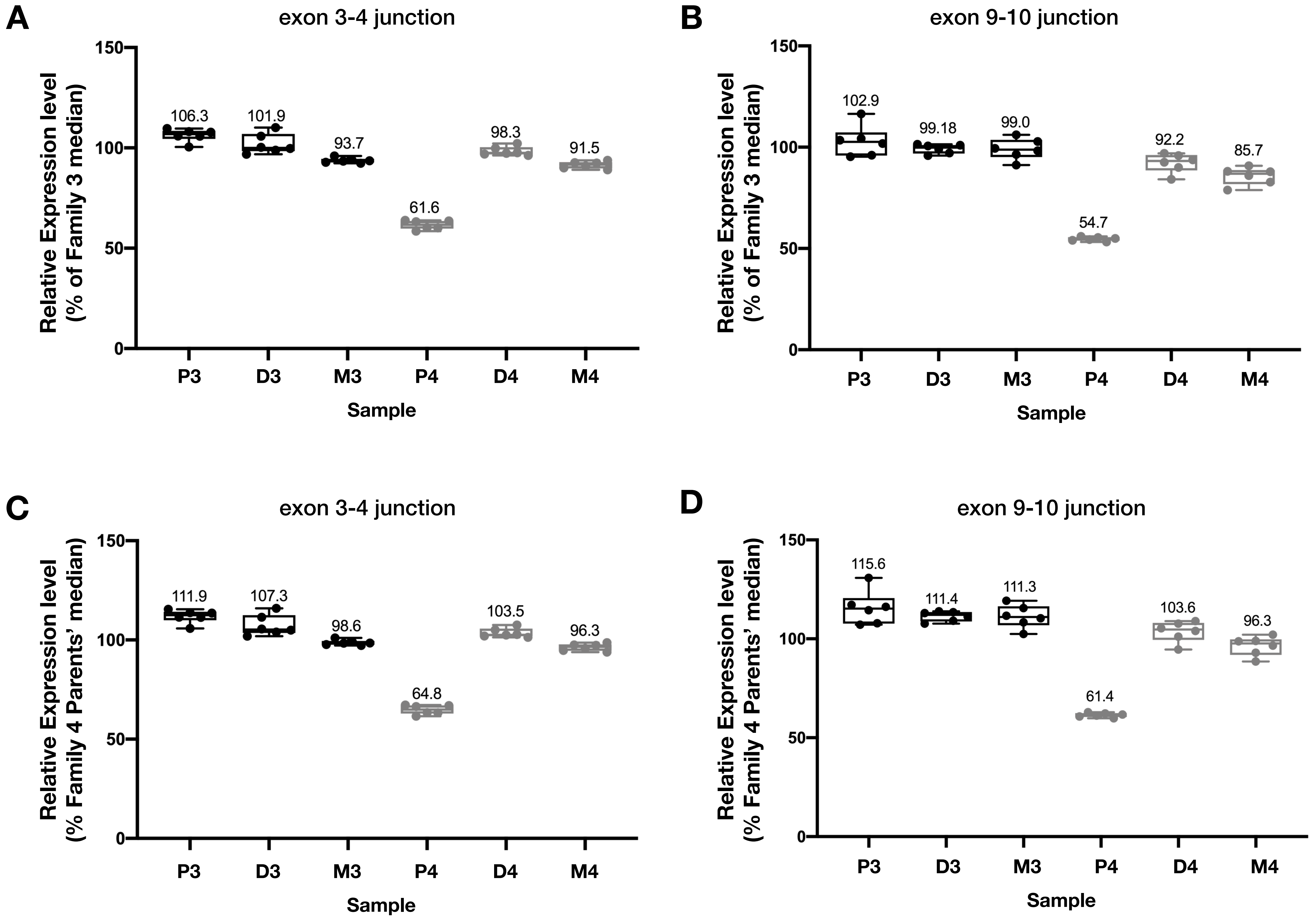


**Supplemental Figure 7. *MLLT3* shows decreased expression in Proband 4 (P4).** qRT-PCR using TaqMan probes targeting the *MLLT3* exon 3-4 (**A, C**) and exon 9-10 (**B, C**) splice junctions, normalized to either the median of Family 3 values (**A, B**), or the median of Family 4 Parent values (**C, D**). Samples include Proband (P), Dad (D), and Mom (M) from two different trios in this study, 3 and 4.

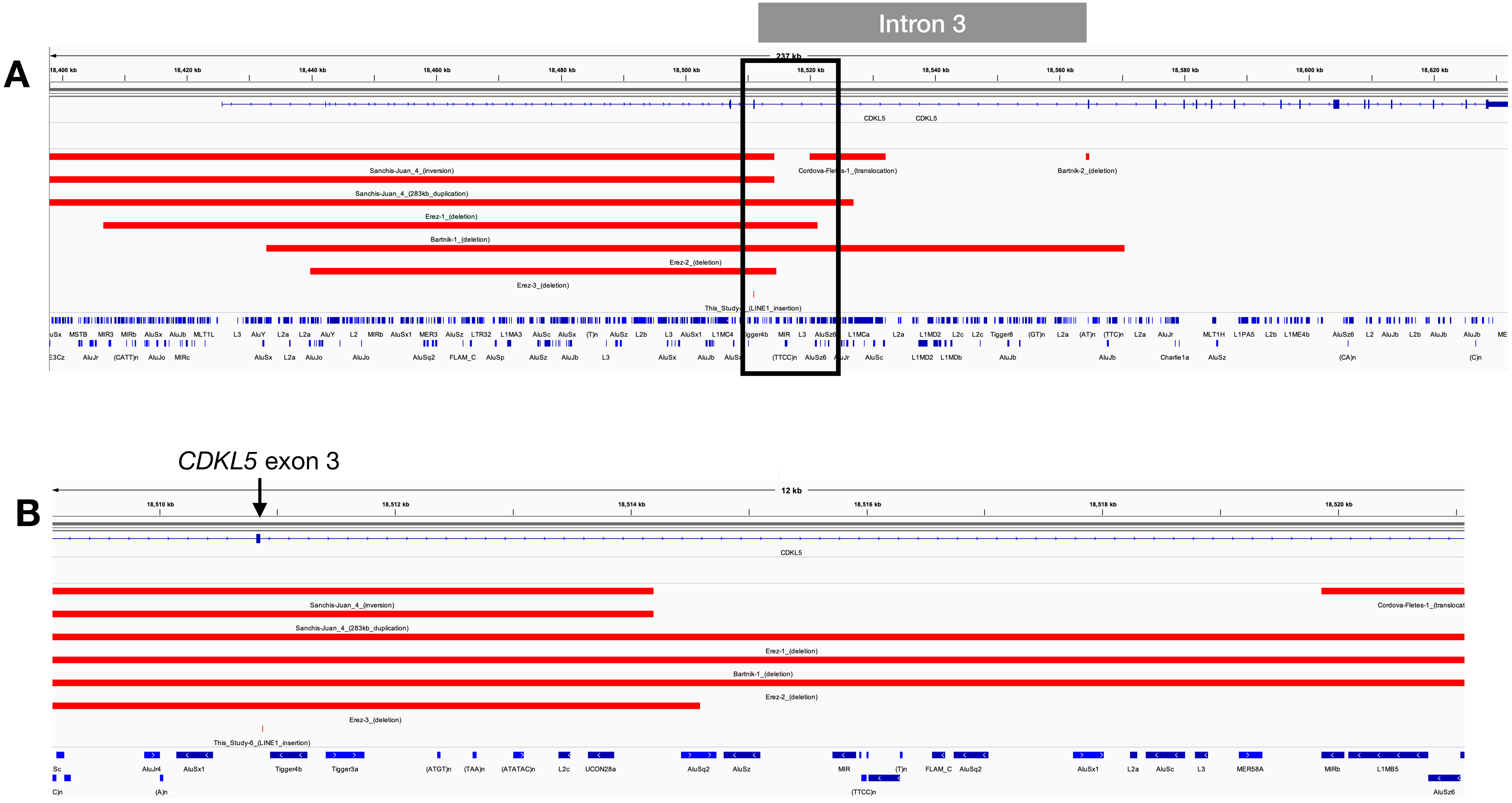


**Supplemental Figure 8.** Several *de novo* structural variants have been reported in *CDKL5*, each with a breakpoint in intron 3 (gray box), near the breakpoint identified here. B is a zoomed in view of A (black box).

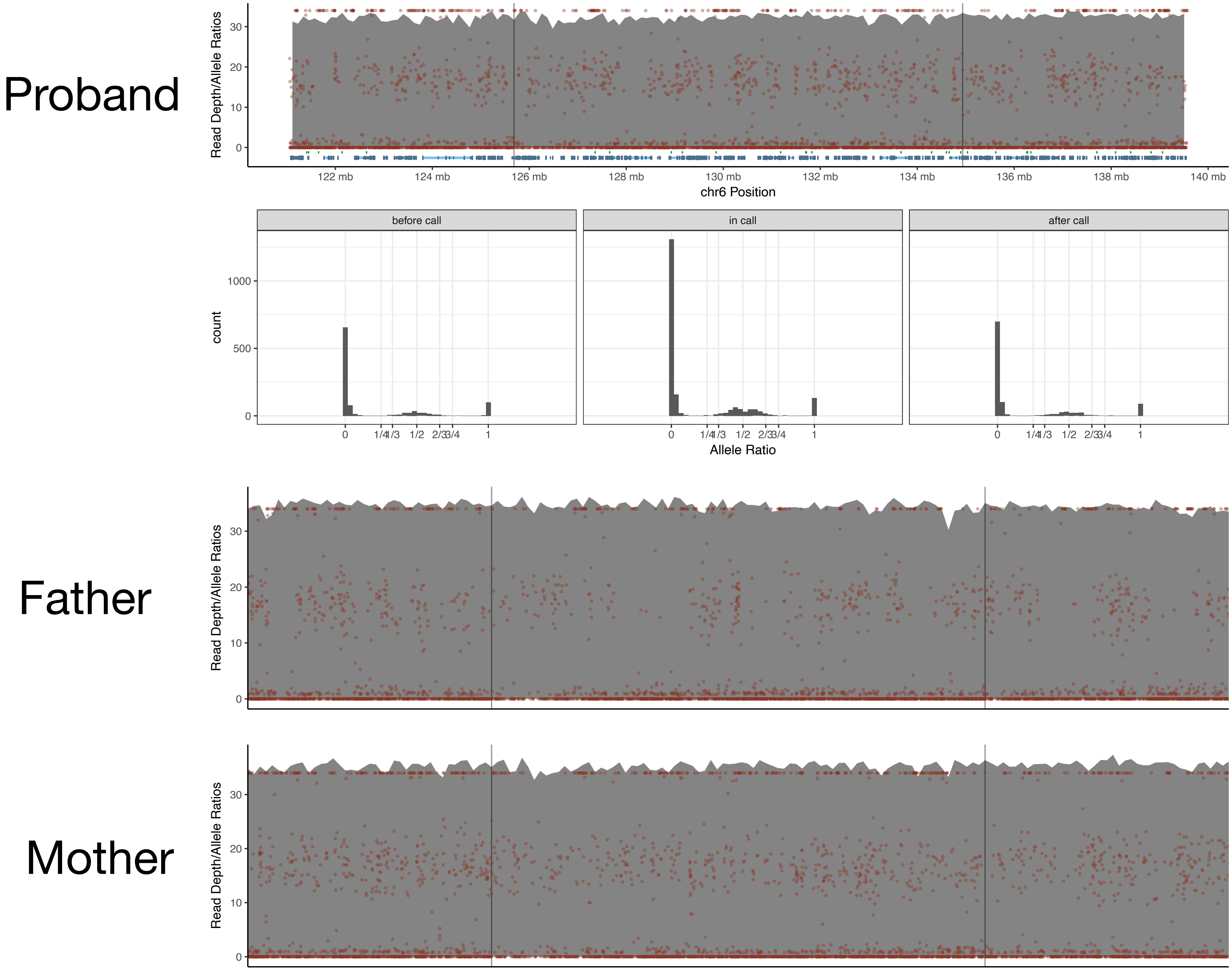


**Supplemental Figure 9.** The complex rearrangements at 6q22.31-6q23.3 appear to be copy-neutral. Plots of read depth (gray peaks) and allele ratios (red dots and plots in second panel from top) in IGS data show similar patterns for proband, mother and father across the region.
